# Canonical retinotopic shifts under an inverse force field explain predictive remapping

**DOI:** 10.1101/2021.09.10.459799

**Authors:** Ifedayo-EmmanuEL Adeyefa-Olasupo, Zixuan Xiao, Anirvan S. Nandy

## Abstract

Predictive remapping — the ability of cells in retinotopic brain areas to transiently exhibit spatio-temporal retinotopic shifts beyond the spatial extent of their classical receptive fields — has been proposed as a primary mechanism that stabilizes our percept of the visual world around the time of saccadic eye movements. Despite the well documented effects of predictive remapping, no study to date has been able to provide a mechanistic account of the neural computations and architecture that actively mediate this ubiquitous phenomenon. We propose a novel neurobiologically inspired general model of predictive remapping in which the underlying pre-saccadic attentional and oculomotor signals manifest as three temporally overlapping forces that act on retinotopic brain areas. These three forces – a centripetal one toward the center of gaze, a convergent one toward the saccade target and a translational one parallel to the saccade trajectory – act in an inverse force field and govern the spatio-temporal dynamics of predictive remapping of population receptive fields. The predictions of our model are borne out by the spatio-temporal changes in sensitivity to probe stimuli in human subjects around the time of an eye movement and are consistent with findings of predictive shifts in the receptive fields of cells in the superior colliculus, frontal eye fields, lateral intraparietal area, and visual area V4.

## INTRODUCTION

In humans, the central 2° of visual space is extensively represented in the retina, superior colliculus, and in the visual cortex and is therefore well suited for everyday tasks which require a detailed inspection of visual objects of interest. During active sensing, saccadic eye-movements are constantly recruited by the visual system to bring objects of interest that fall on the peripheral retina into the central 2° of visual space [1]. Saccades cause large and rapid displacements of the retinal image. These displacements introduce significant disruptions, similar to those observed when attempting to photograph a rapidly moving object using a camera. However, the human visual system has evolved to account for these sudden image disruptions such that our perception of the visual scene remains relatively continuous and stable. The ability of the visual system to account for these disruptions is known as spatial constancy [2-6].

Translational remapping — the ability of cells in the superior colliculus, the frontal eye fields, the lateral intraparietal area, and extrastriate visual areas to predictively shift their sensitivity beyond the spatial extent of their receptive fields (RF) towards their future post-saccadic locations— has been proposed as a primary mechanism that mediates this seamless visual percept [7-11]. However, more recent electrophysiological studies in the frontal eye fields and the extrastriate areas have challenged this dominant view of predictive remapping and have instead shown that receptive fields converge around a peripheral region of interest which includes the location of the saccade target. This form of predictive remapping is referred to as convergent remapping [12-14].

Translational and convergent accounts of remapping thus remain at odds, and the functional role that these divergent forms of transient RF shifts play in mediating spatial constancy remains unresolved. [15-18]. Prior studies have routinely sampled regions in space that could likely yield results favorable to one form of remapping over another [19-20]. In fact, with unbiased spatial sampling, a more recent study suggests that the functional correlates of convergent remapping may include a translational component [21], an account which contradicts an earlier study [16]. Equally unresolved is the issue of time. A recent study has reported that predictive remapping in visual area V4 includes a translational component followed by a convergent component, with each component occurring within distinct non-overlapping temporal windows [22,23]. This temporal account of predictive remapping is however at odds with the canonical order of pre-saccadic and post-saccadic events [6, 24] and a more recent study has challenged this result on methodological grounds [14].

In addition to these unresolved issues, there are three important issues pertaining to the visuo-motor system that have been previously overlooked. First, the frequency of saccades (2-4 times per second) [25] and the accompanied predictive receptive field shifts can be energetically expensive [26]. Yet the neural computations that balance energy costs with predictive function are unknown. Second, the neural architecture that supports neural sensitivity beyond the classic center-surround structure while preventing radical and unsustainable forms of remapping (e.g., over-convergence or over-translation) remains unexplored. Finally, no study has been able to uncover the mechanistic underpinnings that ensure the immediate availability of neural resources at the post-saccadic retinotopic location of the saccade target [27-28]. This rapid availability of neural resources cannot be explained by pure translational shifts towards the retinotopic cell’s future field, or by the spreading of neural resources around the peripheral region of interest.

To provide an account for (a) the spatio-temporal characteristics that define predictive remapping, the neural computations and architecture that ensure appropriate levels of visual sensitivity while preserving retinotopic organization, and (c) the concomitant jump in late pre-saccadic sensitivity in the periphery and the immediate availability of neural resources at the future center of gaze, we systematically assessed the transient consequences of predictive remapping on sensitivity to visual probes along and around the path of a saccade. Next, we investigated the rhythmicity that actively supported these changes across space. Finally, we proposed a novel neurobiologically inspired phenomenological model that succinctly captures the essence of our empirical observations. Additionally, our simulation results (a) align with predictions proposed by classical and more recent neurophysiological and functional studies [7-14, 19, 22-23, 27, 29], (b) explains conflicting neural findings related to whether remapping is limited to the current and future post-saccadic locations [7-12, 27, 29] and (c) provides a set of novel predictions that can inform future investigations.

## RESULTS

### Spatio-temporal profile of pre-saccadic and post-saccadic changes in sensitivity

We assessed changes in visual sensitivity across visual space using a cued saccade task (Fig 1). We specifically examined changes at points along the saccade trajectory (‘radial axis’) and at points orthogonal to the saccade trajectory (‘tangential axis’), while subjects fixated, planned, and executed a saccadic eye-movement toward the saccade target. We probed specific radial points at foveal (*rad*_*fov*−*out*_, *rad*_*fov*−*in*_), parafoveal (*rad*_*para*−*in*_, *rad*_*para*−*out*_) and peripheral (*rad*_*peri*−*in*_, *rad*_*peri*−*out*_) locations, and tangential points symmetric to one another with respect to the radial axis at foveal (*tan*_*fov*−*ccw*_, *tan*_*fov*−*cw*_), parafoveal (*tan*_*para*−*ccw*_, *tan*_*para*−*cw*_) and peripheral (*tan*_*peri*−*ccw*_, *tan*_*peri*−*cw*_) locations, before and after the central movement cue. In each trial, a low contrast probe was flashed at one of these locations (chosen at random) for 20ms with 75% probability; in 25% of trials (control trials) no probe was flashed. The contrast of the probe stimuli was chosen (independently for each subject and for each spatial location cluster; see Methods) such that detection probability was at 50% in the absence of eye movements. The probes were presented at a random time from 600ms before to 360ms after the central movement cue. Our design thus allowed us to measure visual sensitivity along and around the entire saccade trajectory, within early pre-saccadic (−300ms to - 250ms with respect to saccade onset), mid pre-saccadic (−250ms to −150ms), late pre-saccadic (−150ms to - 5ms) and post-saccadic (−5ms to +150ms) windows.

**Figure 1.**
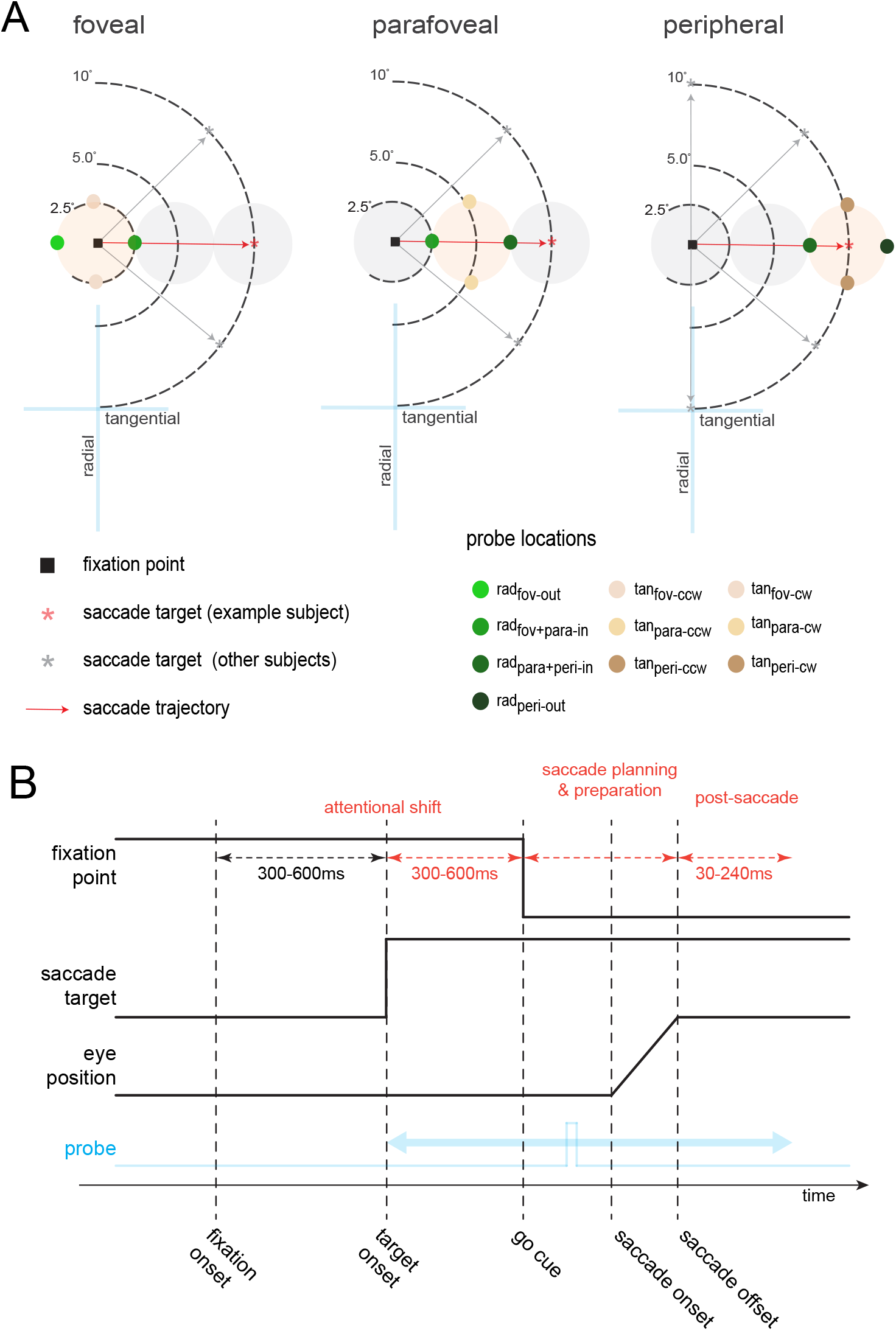
Cued probe-detection task. (A) Spatial locations tested across experiments. Colored circles show the locations at which a low-contrast probe was flashed as subjects planned and prepared a saccade (black arrow) towards the periphery. Probe locations are shown only for horizontal saccades to the right. For other saccade target locations, the probe locations were appropriately positioned along and around the path of a saccade. In the foveal and parafoveal experiments, a saccade target (gray or red asterisk) was presented at azimuth angles of 0°, 45°, 315° (each subject had a different angle). In the peripheral experiment the saccade target was presented either at azimuth angles of 0°, 45°, 90°, 270°, 315° (each subject had a different angle). (B) Temporal sequence of an example trial. The start of a trial began with the appearance of a fixation point, followed by a time-varying presentation of the saccade target. A low-contrast probe was flashed either before the onset of the central movement cue (the pre-saccadic condition), or after (the post-saccadic condition).

Subjects reported whether they were able to detect the flashed probe using a push button. The control (no probe) trials allowed us to assess the incidences of false alarms. Low false alarm rates of 1.2%, 1.4% and 1% (along the radial, tangential counterclockwise and clockwise axes respectively) gave us high confidence about subjects’ visual sensitivity reports. We collected ∼ 21,000 trials, with a minimum of 1,500 trials per subject. Control trials were excluded from our main analyses. We calculated the normalized average visual sensitivity of eleven subjects as a function of flashed probe times relative to saccade onset. Corresponding error estimates were obtained using a 20-fold jackknife procedure in which the sensitivity was estimated from 95% of the data (see Methods).

Within an early pre-saccadic window (−300ms to −250ms) we found a graded profile of sensitivity change along the radial axis [29]: an immediate and sharp decline in visual sensitivity in the outer peripheral region (*rad*_*peri*−*out*_), a modest decline in visual sensitivity at the outer parafoveal region (*rad*_*para*+*peri*−*in*_), and sustained levels of visual sensitivity in the foveal (*rad*_*fov*−*out*_) and inner parafoveal (*rad*_*fov*+*para*−*in*_) regions. (Fig.2, left panel). Functionally, this is in alignment with prior reports of pre-saccadic compression of visual space toward the fovea as subjects maintain fixation just before the deployment of attention towards the periphery [30-32]. This was followed by a modest rebound in sensitivity within a mid pre-saccadic window (−250ms to −150ms) at the more eccentric locations (*rad*_*peri*−*out*_ and *rad*_*para*+*peri*−*in*_) from 240ms to 130ms prior to saccade onset, consistent with a pre-saccadic shift of attention toward the saccade target [6, 12-14, 16]. Within a late pre saccadic window (−150ms to −5ms) just before saccade onset, we found continued declines in sensitivity at locations near the current center of gaze (*rad*_*fov*−*out*_, *rad*_*fov*+*para*−*in*_). Aligned with classical neurophysiological findings [7-11, 27], we found a concomitant jump in late pre--saccadic sensitivity first at the peripheral location radially outward from the saccade target (*rad*_*peri*−*out*_) followed immediately by a similar jump at the less peripheral radially inward location (*rad*_*para*+*peri*−*in*_). Finally, within a post-saccadic window (−5ms to +150ms) we found continual declines in post-saccadic sensitivity, as we had expected, in the fovea and inner parafoveal regions. This global decline in sensitivity is in agreement with late pre--saccadic corollary discharge commands originating from pre-motor brain areas as well as with visual-transient signals originating from the retina to retinotopic brain areas just before subjects executed a saccade [33-36]. Concurrently, there was a rapid increase in post-saccadic sensitivity at the locations around the saccade target (*rad*_*peri*−*out*_, *rad*_*para*+*peri*−*in*_) [28].

**Figure 2.**
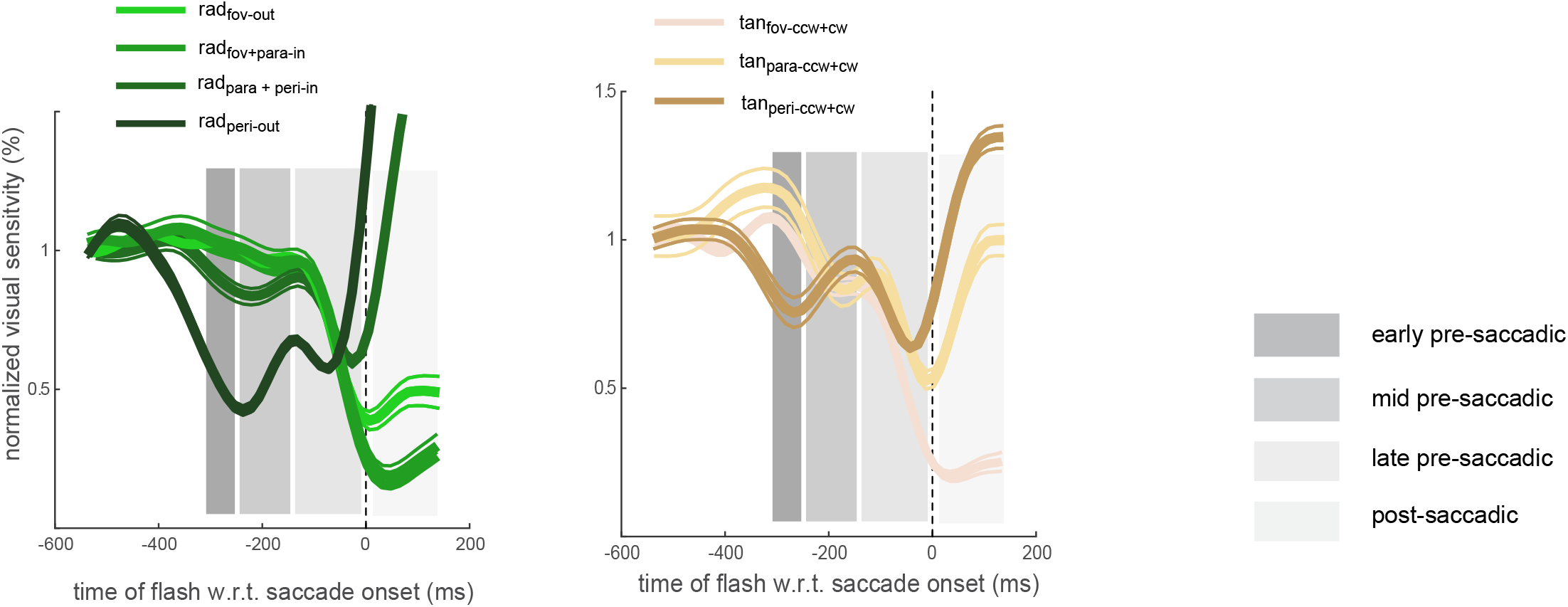
Sensitivity functions across visual space within distinct pre and post saccadic windows.Thick solid lines represent normalized changes in pre and post-saccadic sensitivity as a function of flashed probe times relative to saccade onset along and around the saccade trajectory (n=11). The thin lines represent corresponding error estimates obtained using a 20-fold jackknife procedure in which the sensitivity was estimated from 95% of the data.

Our observations of sensitivity changes along the tangential axes largely mirrored those along the radial axis in an eccentricity dependent fashion (Fig 2, right panel). We found a decrease in sensitivity at the *tan*_*peri*−*ccw*+*cw*_ location within early pre-saccadic window. This was accompanied by a modest increase in mid pre-saccadic sensitivity in the parafoveal region (*tan*_*para*−*ccw*+*cw*_) or sustained levels of sensitivity in the foveal (*tan*_*fov*−*ccw*+*cw*_) region. Similar to the radial axis, we observed a rebound in late pre-saccadic sensitivity (−50ms from saccade onset) at the most eccentric location (*tan*_*peri*−*ccw*+*cw*_) that was followed by a similar rebound at the parafoveal location (*tan*_*para*−*ccw*+*cw*_). A continued decline in post-saccadic sensitivity at the foveal location was accompanied by rapid increases first at the peripheral locations followed by the parafoveal location.

### Low-frequency rhythmicity underlies changes in sensitivity

Our behavioral results suggest that the changes in pre- and post-saccadic sensitivity are much more dynamic than previously reported [15-18]. Specifically, our results suggest that neural resources are initially allocated towards less eccentric retinotopic locations, allowing for a graded profile of sensitivity across visual space in favor of the current center of gaze. After transient shifts in sensitivity toward the saccade target however, these graded changes in visual sensitivity disappear as the more eccentric locations experience quicker increases in late pre-saccadic sensitivity well before observed increases at intermediate locations (Fig 2). We hypothesized that these dynamics are a consequence of attentional shifts during fixation and around the time of a saccade. To test this, we assessed the spectral signature of the observed changes in sensitivity [37]. The logic behind this assessment is as follows: if these changes are a consequence of attentional shifts, then the spectral signature associated with these changes should resemble, if not match, known rhythmic patterns of attentional shifts during fixation and around the time of a saccade. Conversely, if the spectral architecture we find fails to recapitulate these known rhythmic patterns, then these changes cannot be directly attributed to attentional shifts.

Visual sensitivity around the current center of gaze (“foveal”) locations (*rad*_*fov*−*out*_, *rad*_*fov*+*para*−*in*_) exhibited rhythmicity within a delta spectral window at ∼2 Hz, with the exception of the *tan*_*fov*−*ccw+cw*_ location which exhibited rhythmicity that bordered the delta and theta windows at ∼3 Hz (Fig 3A, left panel). Spectral differences calculated using location shuffled data (n=1500) versus the non-shuffled data at these locations were statistically significant (two-tailed paired-sample t-test, p=0.0018, p=0.0236, p=0.0018) (Fig 3A, right panel). At the intermediate (“parafoveal”) locations (*tan*_*para*−*ccw*+*cw*_), we found that sensitivity exhibited delta-band rhythmicity at ∼2Hz (Fig 3B, left panel), but spectral differences between the shuffled and non-shuffled data sets were not significant (p=0.1738). However, peripheral locations around the future center of gaze (*tan*_*peri*−*ccw*+*cw*_, *rad*_*para*+*peri*−*in*_, *rad*_*peri*−*out*_) all exhibited rhythmicity at 2-3Hz which were statistically significantly different from location-shuffled data (p= 0.0062, p=0.0071, p= 0.0207 respectively). Taken together, our results suggest the presence of location-specific rhythmic sensitivity in task-relevant regions of visual space around the current and future centers of gaze [6, 19, 34, 35]. In line with our hypothesis, the frequency band at ∼2Hz is known to support pre-saccadic attentional shifts that mediate visual sensitivity at suprathreshold levels [38-40]. Additionally, the frequency band at ∼3Hz predicts attentional shifts associated with small fixational eye movements [41].

**Figure 3.**
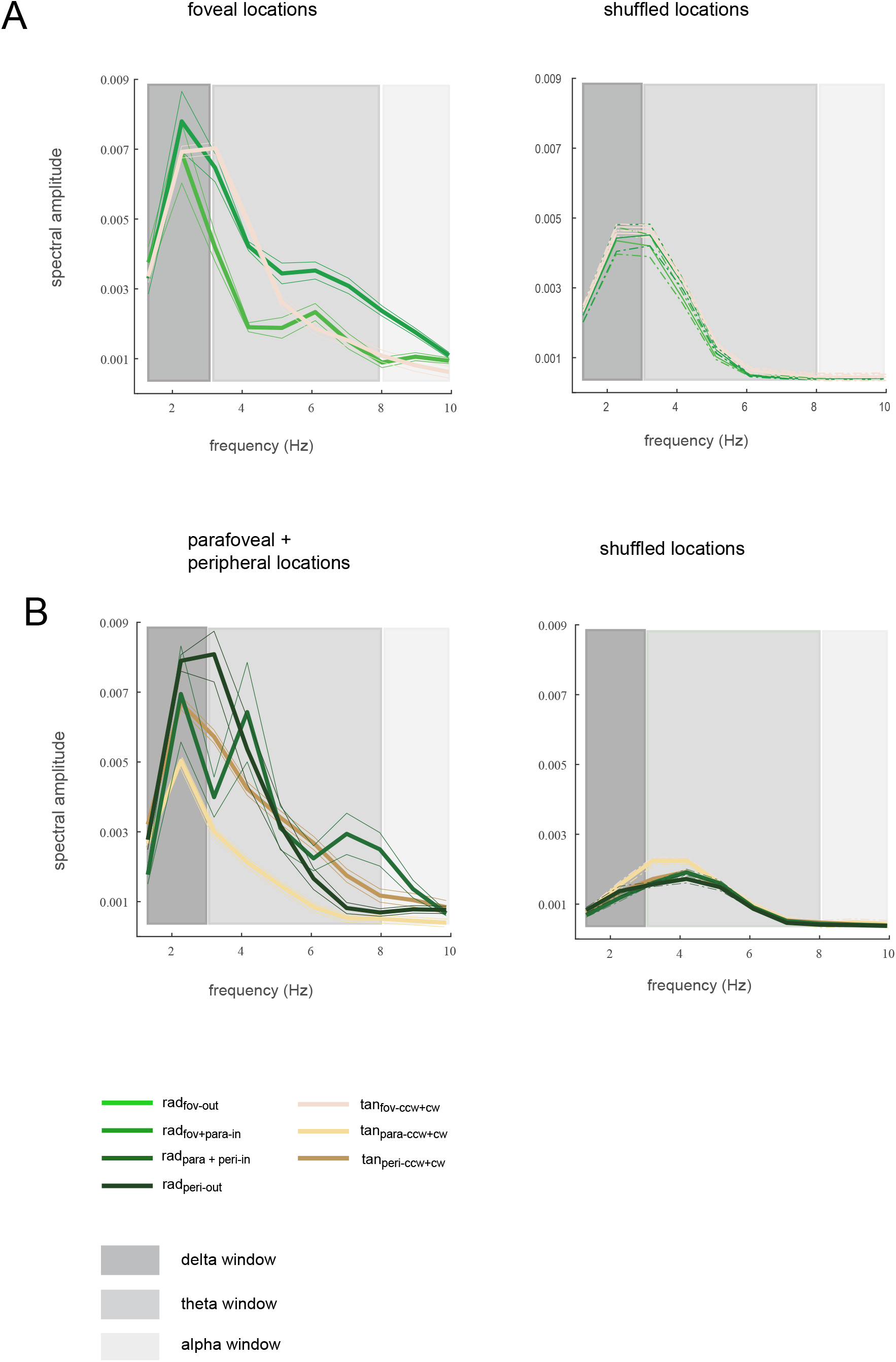
Frequency domain representation of detrended sensitivity functions within distinct spectral windows. (A) Frequency domain representation of detrended visual sensitivity, unshuffled and shuffled (n=1500) in the fovea. The thinner solid lines represent the error estimates calculated across subjects. (B) Unshuffled and shuffled (n=1500) in the more eccentric regions.

### Force field model of predictive remapping

Population receptive fields (pRFs) across retinotopic brain areas are not static but can exhibit transient spatial shifts [7-14]. To this end, we developed a simple yet powerful phenomenological model that allows the pRFs of cells in retinotopic brain areas to transiently undergo spatio-temporal retinotopic shifts beyond the spatial extent of their classical receptive fields consistent with translational, convergent as well as other biologically plausible forms of remapping.

Our model assumes that there are limits to the extent of these transient retinotopic shifts such that retinotopic organization is always maintained [42]. Consequently, within our model we extend the idea of a neuron’s receptive field by introducing the concept of its elastic field. We posit that population elastic fields (pE∅) are self-generated by the visual system and constitute the spatial limits within which pRFs are allowed to transiently shift. These transient spatial shifts manifest as distortions of pRF density [11, 13, 31, 42] and temporarily distort the representation of visual space [43-44]. pE∅ span the region immediately beyond the classical extent of the pRF, their spatial extent is proportional to the eccentricity of the classical pRF, and they are omnidirectional with respect to the classical pRF (Fig 4A).

**Figure 4.**
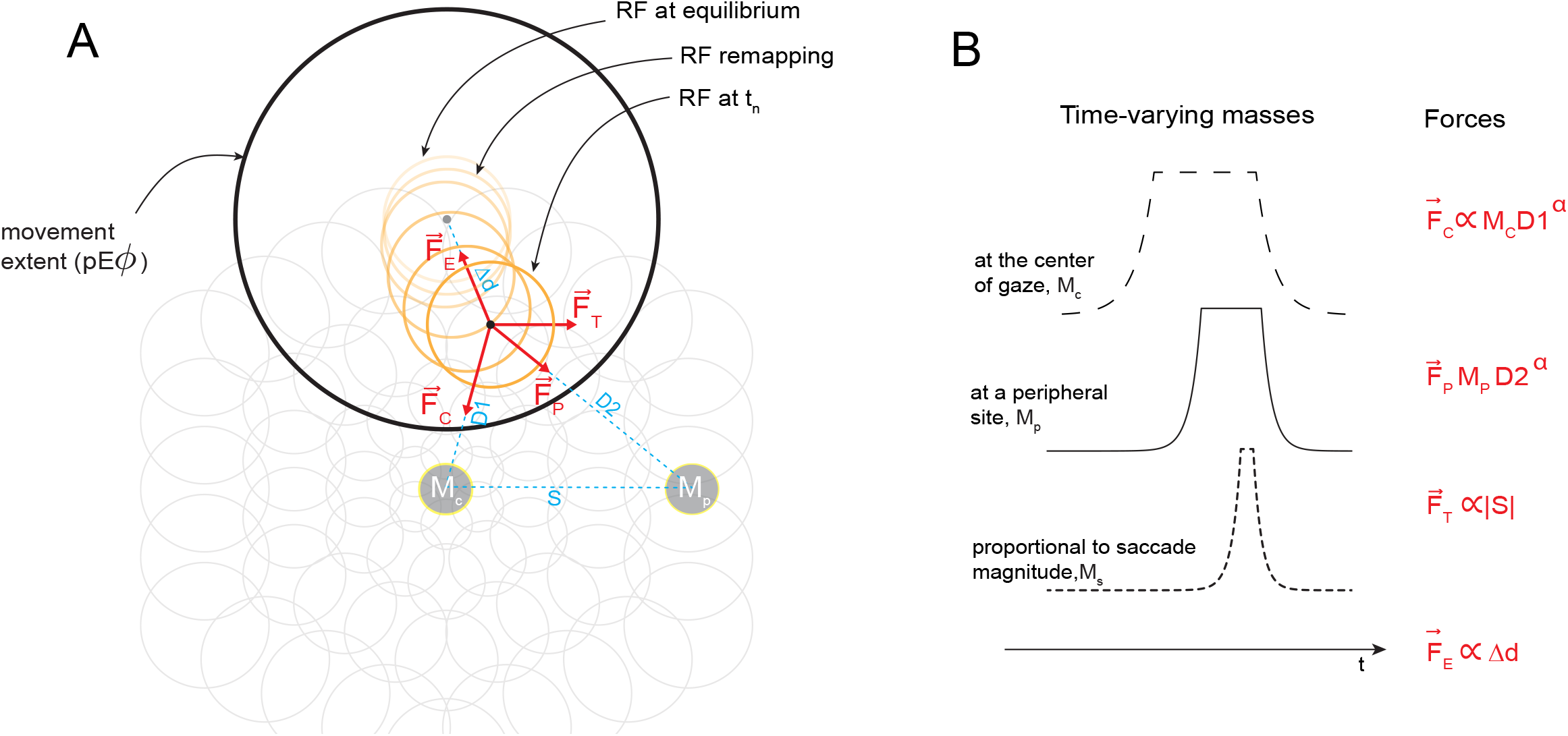
Neural architecture and mechanics underlying the force field model of predictive remapping. Retinotopic field consisting of population receptive field (pRFs) that tile visual space. A single pRF is highlighted in orange. This pRF is characterized by the location of its center at equilibrium (gray dot), its size (orange circle) and a movement extent (elastic field, pE black annulus). Here movement extent of pRF is exaggerated for the sake of or illustration purposes. (B) Both pRF size are eccentricity dependent. M_C_ represents centripetal signals and is modelled as a varying mass located at the center of gaze. M_P_ represents convergent signals and is modeled as a varying mass located at the peripheral site corresponding to the saccade target. M_T_ represents translational signals and is modelled as a virtual mass at infinity located in the direction of the impending saccade.

Visually transient signals originating from the retina and corollary discharge signals originating from pre-motor brain structures are sent to relevant retinotopic brain areas [7-14, 27-33, 36, 45-46]. We will refer to these signals, in aggregate, as “attentional-oculomotor” signals. At its core, our model assumes that attentional-oculomotor signals act as forces that perturb pRFs from their equilibrium positions. We model these forces as being exerted by corresponding time varying masses (M) in a “retinotopic force field”. The perturbations due to these forces are limited to the spatial extent of pE∅ and are inversely proportional to the distance between the pRF and M (Fig 4A,B). These principles ensure that pRFs are appropriately sensitized around the loci of task-relevant salient cues.

Prior to a saccadic eye-movement, attentional-oculomotor signals are thought to arrive earlier at the fovea but are biased in the direction of the saccade target [31, 46, 47]. Therefore, both the current center of gaze and the path to the saccade target inherently benefit from earlier and faster visual processing, reflecting a detailed visual representation within these retinotopic regions [30]. Conversely, at the more eccentric parts of the visual field, visual processing is slower and yields a more diffuse visual representation within these retinotopic regions [12, 44]. Considering these previous results, we posit that pre-saccadic attentional and oculomotor processes manifest in three forces that aligns with the spatio-temporal attributes of canonical pre-saccadic events [6, 24, 46 48] and impinge on the retinotopic visual cortex (Fig 4B). We hypothesize that the transient perturbations of pRFs caused by these forces are the underlying causes of predictive remapping (Fig 4B).

The earliest force – a centripetal one, overlapping with the time of fixation – causes pRFs to transiently exhibit centripetal shifts towards the current center of gaze. We hypothesize that these centripetal shifts facilitate the acquisition and storage of information and resources the visual system will eventually transfer towards the future center of gaze [29, 49-50]. The centripetal force is followed by a convergent force, which aligns with the deployment of pre-saccadic attention towards the peripheral target and causes pRFs to transiently exhibit convergent shifts around the peripheral site [12-14]. This convergence provides a perceptual advantage around the task-relevant target [6, 21]. Finally, as the organism prepares and plans to execute the imminent saccadic eye-movement, a translational force causes pRFs to shift towards their future receptive fields [7-12]. This, in combination with a declining centripetal force (which in our model represents the saccade onset), ensure a handoff of neural resources toward the future center of gaze [28].

### Systematic investigations and novel predictions associated with the different components of predictive remapping

The underlying architecture of our model consists of two main layers – a “retinotopic field” and a “force field”. The retinotopic field is composed of overlapping pRFs that tile visual space. The size of each pRF is proportional to its eccentricity, as is the corresponding elastic field (pE∅). Force fields perturb pRFs from their equilibrium positions. This field is activated by the availability of attentional-oculomotor signals (“forces”) introduced within canonical overlapping temporal windows (Fig 4B). Each force is exerted by its corresponding mass (M) which is modelled with the following distributive parts: an exponential growth, a stable plateau, and an exponential decay respectively. The perturbation of pRFs produces spatiotemporal retinotopic shifts constrained by their pE∅s (Fig 5) which manifest in time-varying modulation of neural population density readouts. We assume that these neural density readouts are equivalent to stimulus detection sensitivity at the psychophysical level.

**Figure 5.**
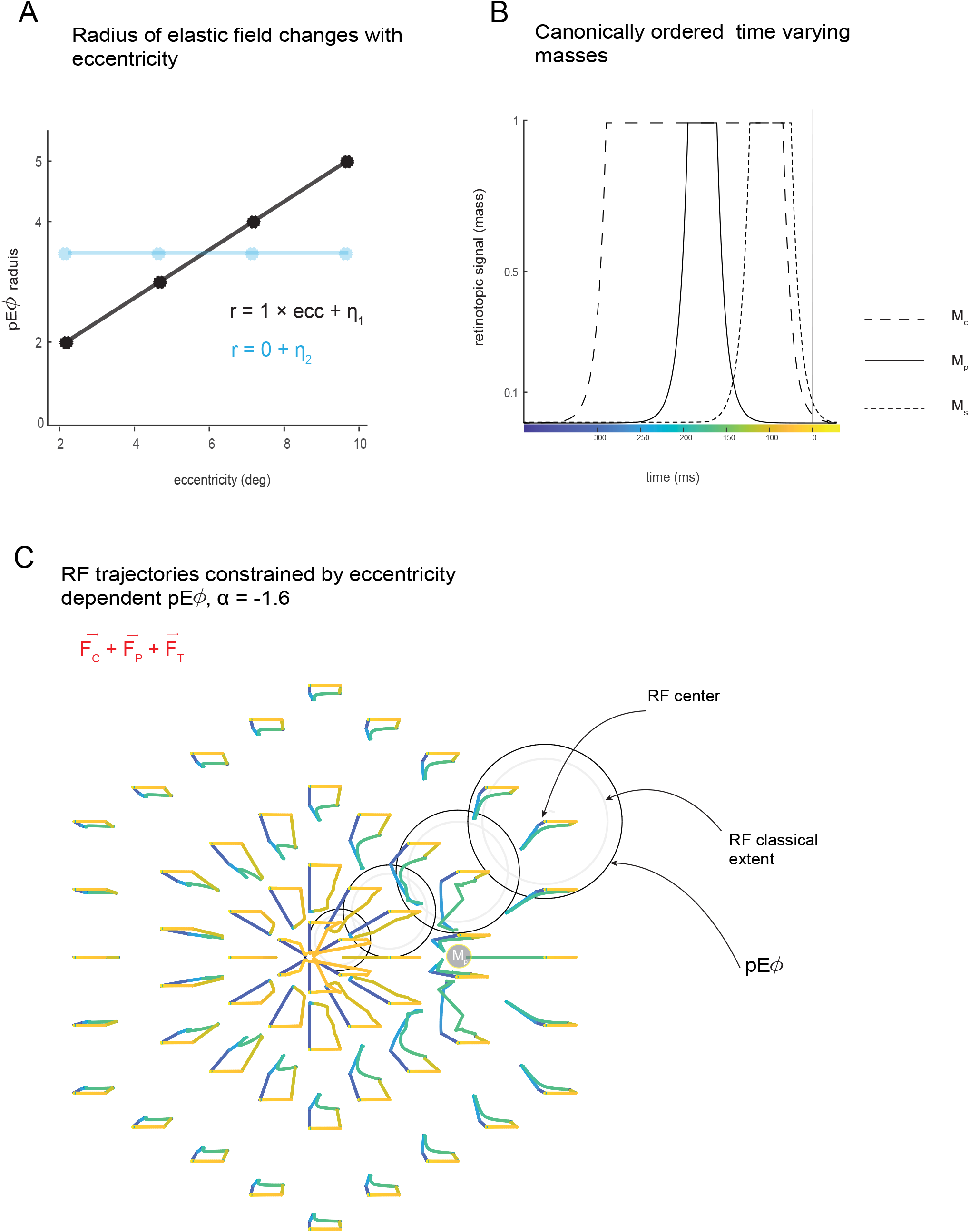
Retinotopic shifts under an inverse force field driven by simulated centripetal, convergent and translational retinotopic signals. (A) population elastic fields (pe *ϕ*) can either be: eccentricity dependent (black trace), non-eccentricity dependent (blue trace). In addition to these two dynamics, there is third case where pRFs do not possess elastic fields during active vision (B) time varying masses for the case when pRFs were perturbed the linear combination of three types of retinotopic signals modelled as independent forces. (C) Corresponding spatiotemporal pRF trajectories under an inverse force field (α = −1.6). Note that each pRF possess an eccentricity dependent pEF, however for illustration purposes only a few are shown (black circles). The trajectories are color coded as in (B).

To systemically investigate the contributions of different model components as well as their interactions within specific temporal windows, we performed simulations with various combinations of forces (Fig 6-7), under an inverse force field where *α* is set to −1.6. We report neural density readouts at locations along the radial axis that corresponded to those sampled in our psychophysical experiment: outer foveal, inner foveal/inner parafoveal, outer parafoveal/inner peripheral, and outer peripheral. These sampled retinotopic locations allowed us to assess the generalizability of our model in the foveal, parafoveal and peripheral regions of visual space.

**Figure 6.**
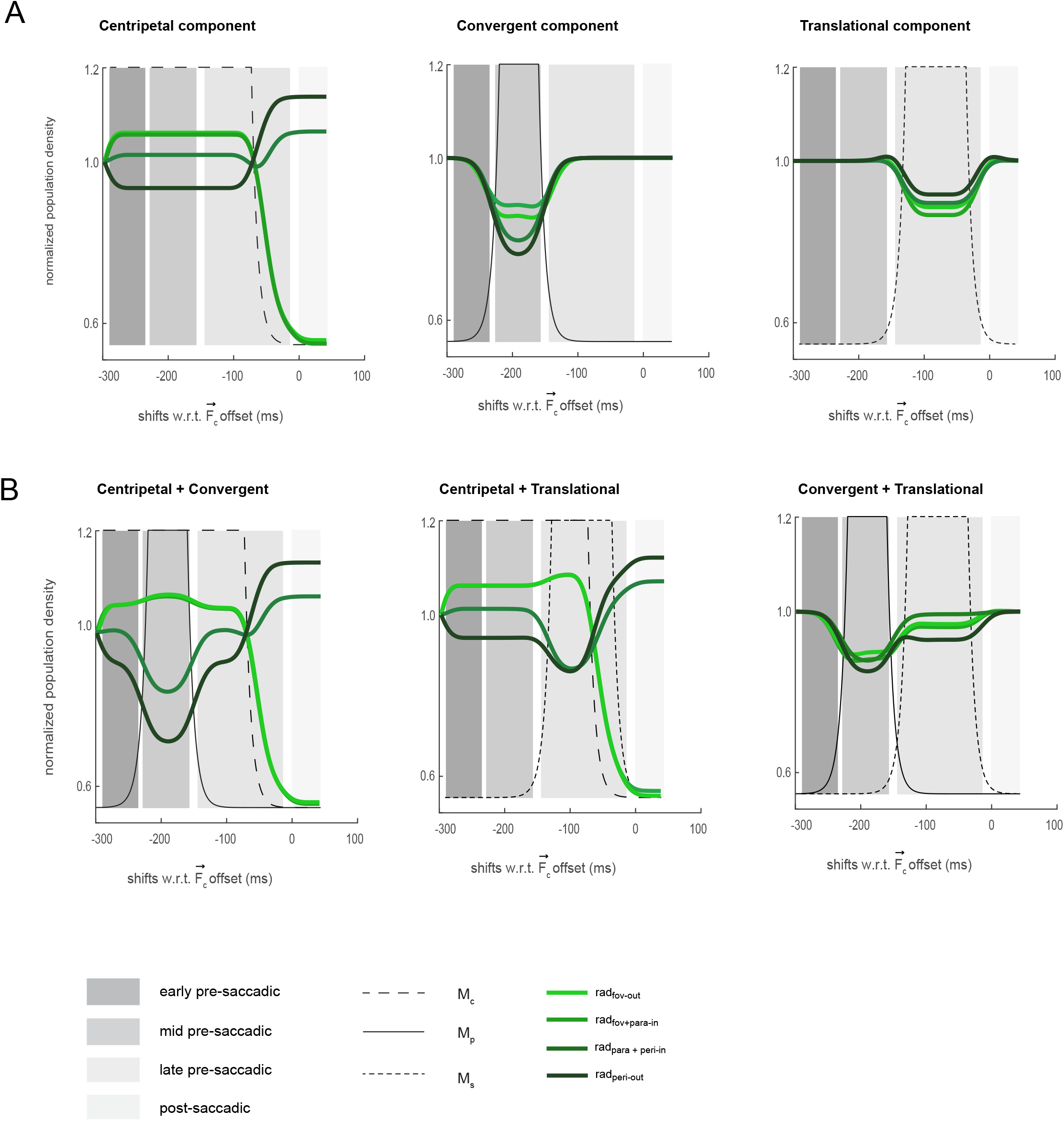
Simulated population neural signatures under an inverse force field. (A) Simulation results at α = −1.6 in cases when pRFs are perturbed by a single external force constrained by eccentricity dependent pE*ϕ*s. Along x-axis represents the simulation time sampled at discrete increments of 1ms realigned with respect to the offset of the central force (i.e., modelled as the saccade onset) (B) when pRFs are perturbed by the summation of two external forces constrained by eccentricity dependent pE*ϕ*s.

#### Centripetal component

When modelled pRFs constrained by their pE∅s were perturbed by only a centripetal force, we found distinguishable differences in population density levels between more foveal (*rad*_*fov-out*_ and *rad*_*fov-in*_) and more eccentric (*rad*_*para+peri-in*_ and *rad*_*peri-out*_) locations during the sustained period of this force, (Fig 6A, left panel). The decline of the centripetal force was accompanied by a clear hand-off in population density levels between the foveal and peripheral locations. We found an earlier and stronger increase in population density at the outer peripheral location (*rad*_*peri*−*out*_), followed by an increase in density levels at the inner peripheral location (*rad*_*para*+*peri*−*in*_). These model dynamics mirror our empirical findings during the late pre-saccadic period (Fig 2, left panel) and can be explained as follows: The decline of the centripetal force restores the pRFs to their equilibrium position. The larger size and coverage of the more eccentric pRFs help restore density earlier in the more peripheral location compared to the less peripheral location.

#### Convergent component

Under the perturbation by a convergent force alone, 250ms to 150ms before saccade onset, we found a graded decline in population density levels across visual space (Fig 6A, middle panel). Under a strong inverse force field, the onset of a convergent force causes pRFs that support rad_para+peri-in_ and rad_peri-out_ locations to allocate more of their resources towards the target, while at the more distal locations with respect to the target (*rad*_*fov-out*_ and *rad*_*fov-in*_) fewer resources from these regions are being allocated towards the target. This result aligns with a more recent functional study [21] and a growing number of neural studies, all of which suggest that functional and neural correlates of convergent remapping, which leads to a perceptual advantage around the task-relevant target, fundamentally requires an additional component [12, 14, 22, 23].

#### Translational component

Similar to the simulations for the purely convergent case, we found a graded decline in population density levels across visual space due to a pure translational force applied during a late pre-saccadic window (Fig 6A, right panel). However, here, changes in population density levels favor the future center of gaze over the current center of gaze. These results align with the predictions proposed by early neurophysiological studies that the anticipatory transferring of resources from a cell’s current field to its future post-saccadic location is in part driven by oculomotor related translational signals. On a functional level, this causes a relatively equal and transient decline in sensitivity across visual space [33-35]. It also aligns with a previous study which suggest that the translational signal plays a role in mediating post-saccadic resources [15].

#### Centripetal and convergent components

The combination of centripetal and convergent forces (Fig 6B, left panel) resulted in an amplification of the differences in population density levels along the radial axis during the application of the convergent force (compare with individual force simulations in Fig 6A, middle and left panels), while preserving the handoff noted earlier for the centripetal only simulation. This amplification mirrors our empirical finding during the mid pre-saccadic period (Fig 2A, left panel). Together with the centripetal simulation, these results predict the importance of the centripetal force in not only maintaining the appropriate level of resources at the current center of gaze during the sustained period of the force but also ensuring the immediate availability of neural resources at the future center of gaze during the offset of the force (Fig 6A left panel Fig 6B, left panel) [28].

#### Centripetal and translational components

The combination of the centripetal and translational forces resulted in an amplification of the divergence in sensitivity between the foveal and peripheral locations during the application of the translational force, while retaining the foveal-peripheral handoff during the decline of the centripetal force. This amplification mirrors our empirical findings during the late pre-saccadic period (Fig 2A, left panel).

#### Convergent and translational components

The combination of convergent and translational forces resulted in a noticeable increase in population density levels at the *rad*_*para+peri-in*_ and *rad*_*peri-out*_ locations (Fig 6B, right panel) when compared to the simulation by a convergent force alone (Fig 6A, middle panel). In the neural domain, this change in population density levels around the peripheral target align with previous studies that have shown pre-saccadic transient increases in neural sensitivity in visual area V4 and the frontal eye fields around the saccade target [12-14]. In the functional domain, these changes would constitute a perceptual advantage around the task-relevant target [6, 21]. These results provide the long sought-after computational evidence that translational signals in combination with convergent signals can support transient increases in sensitivity around the target of the impending saccade [12, 13, 14, 20, 21, 22, 23, 44].

#### All components

While the single and paired force simulation capture specific components of our empirical findings and those from the extant literature, the combined effects of all three forces (Fig 7A, left panel) best capture our empirical results (Fig 2A, left panel), suggesting the importance of each of these forces in mediating pre-saccadic sensitivity: the centripetal force is key to maintaining sustained levels of resources at the current center of gaze while ensuring a quick pre-saccadic handoff to the future center of gaze during its offset; the combination of convergent and translational forces support transient pre-saccadic increases in sensitivity around the saccade target.

**Figure 7.**
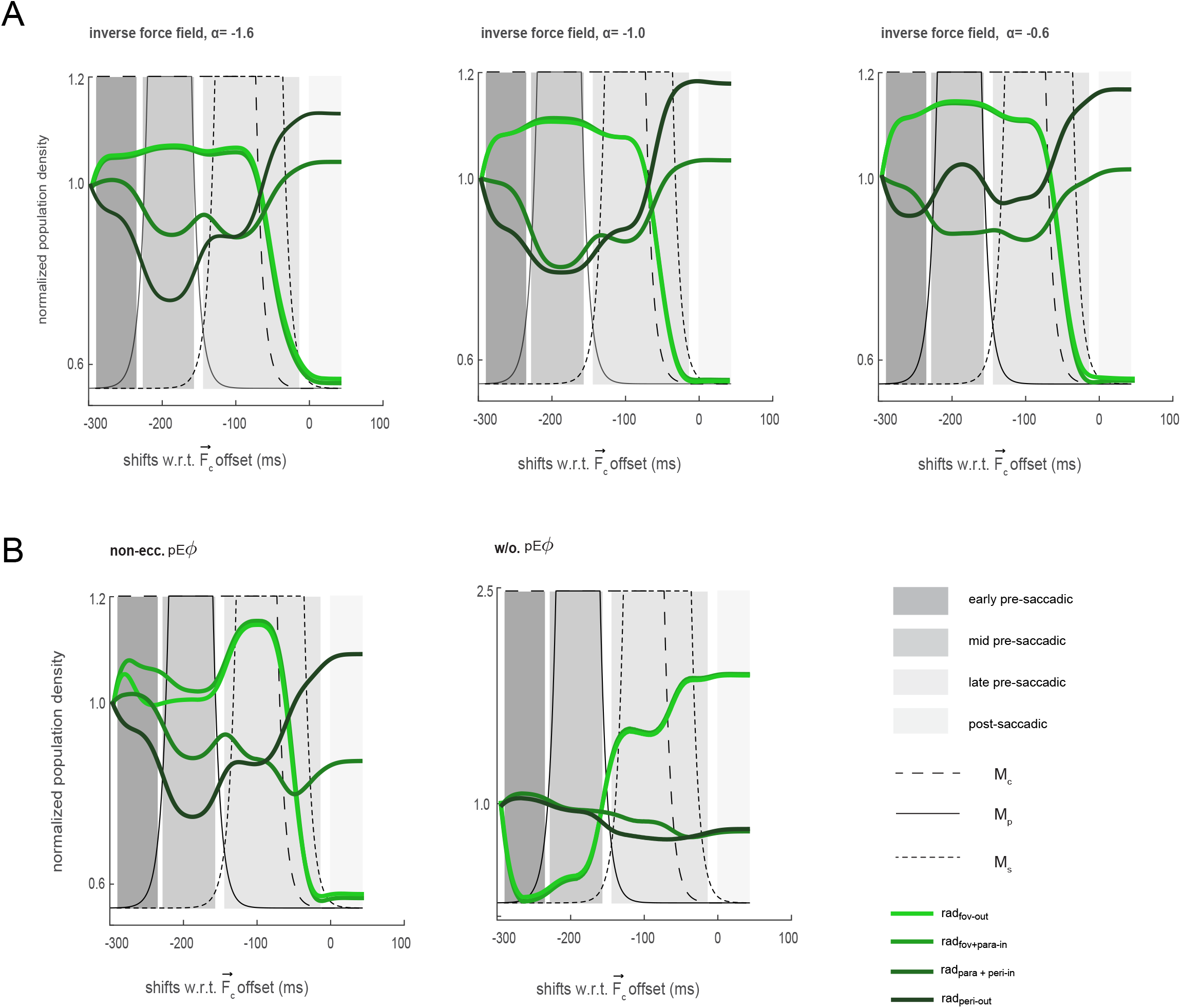
centripetal, convergent and translational signals under an inverse force field explain remapping. (A) Simulated population signatures constrained by eccentricity dependent pE*ϕ*s, under distinct inverse force fields of varying strengths. Along x-axis represents the simulation time sampled at discrete increments of 1ms. (B) Simulated population signatures for the combination of three independent forces that are constrained by non-eccentricity dependent pE*ϕ*s and unconstrained.

Classical neurophysiological studies have shown that retinotopic cells respond to a salient cue placed within their future RF before the onset of an eye movement [7-12, 27]. However, more recent results by Wang and colleagues suggest that cells in LIP also become responsive to intermediate locations along the saccade trajectory [29]. This is said to drive activity in favor of the current center of gaze that is eventually redirected (or remapped) towards the cell’s future post-saccadic location Our simulation results, along with our empirical observations, strongly suggest that graded changes in sensitivity towards the fovea occurs within an early to mid pre-saccadic window, as predicted by Wang and colleagues. However, these graded changes in sensitivity cease within a late pre-saccadic window just before saccade onset as was predicted by more classical studies [7-12, 27]. This is initiated by a declining centripetal force which causes stronger and rapid increases in sensitivity and density levels at the *rad*_*peri*−*out*_ location before any other location in the periphery.

#### Different inverse force fields

To assess the impact of the spatial profile of the force fields, we repeated the three-force simulations with different values of the α parameter. With *α* = −1 (Fig 7A, middle panel), population density signatures at the more foveal locations are largely unchanged when compared to the results under an inverse force field where *α* is set to −1.6 (Fig 7A, left panel). More eccentric pRFs are sensitized towards the current center of gaze but less so when compared to a strong inverse force field. To this end, within an early-mid pre-saccadic window we found declines in population density levels at the *rad*_*para+peri-in*_ and *rad*_*peri-out*_ locations. However, declines at the *rad*_*peri-out*_ location were not as large when compared to what we observed under a stronger inverse force field. Under an inverse force field where *α* is set to −0.6, intermediate parafoveal pRFs are more sensitized towards the current of gaze. To this end, we observed larger declines in population density levels at the *rad*_*para+peri-in*_ location when compared to the simulations under an inverse force field where *α* was set to −1 or −1.6.

#### Population elastic fields

All the results we have presented thus far were simulated with modelled pRFs constrained by eccentricity dependent pE∅s (Fig 4A, black trace). To assess the role of pE∅s in pre-saccadic density dynamics, we ran two additional simulations (both with a strong inverse force field, *α* = −1.6). For the first stimulation, pE∅s were the same size regardless of the eccentricity of the corresponding pRFs (Fig 4A, blue trace), while in the second stimulation pRFs were unconstrained by their presumed pE∅s. In the case of an eccentricity independent tilling of pE∅s, we found no appreciable differences in density levels when compared to the signatures we observed under eccentricity dependent constraints during the early-mid pre-saccadic periods (Fig 7B, left panel; compare with Fig 7A, left panel). However, a different signature emerges with the introduction of a translational force. Specifically, we found that sustained density levels at the foveal regions increase in magnitude, while there is a marked decrease in density level at the more intermediate eccentric location. Eccentricity dependent pE∅s allows more eccentric pRFs a greater movement extent (i.e., more elasticity). Under a non-eccentricity dependent configuration, however, eccentric pRFs are much more restricted in their ability to shift within the visual field. Consequently, once eccentric pRFs become sensitized towards the current center of gaze, they become restricted from reallocating these resources back towards the periphery. This results in an inappropriate level of sensitivity at the current center of gaze within a late pre-saccadic window as opposed locations around the future center of gaze.

Finally, for simulations where pRFs were unconstrainted by their pE∅s, retinotopic organization simply collapses (Fig 7B, right panel). The centripetal force results in an over compression towards the current center of gaze. Further, despite the onset of the convergent and translational forces, pRFs are desensitized to the presence of these forces. This result points to the importance of pEFs in preventing any radical and unsustainable forms of remapping and thus maintaining retinotopic organization.

## SUMMARY

Our study was motivated by what is arguably the most puzzling question in the field of predictive remapping research. In 1990 Goldberg and Bruce recapitulated translational effects observed in the superior colliculus by Mays and Sparks ten years earlier [51-52]. They found that before the execution of a saccade, the receptive fields of cells in the frontal eye fields predictively shift their spatial extent beyond their classical extent towards their future fields (translational remapping). These neural effects were reproduced across several retinotopic brain areas including the lateral intraparietal area and extrastriate visual areas. About two decades later, recapitulating in part the results of Tolias and colleagues [12], Zirnsak et al. demonstrated that the receptive fields of cells in the frontal eye fields primarily converge around the attention-selected peripheral region of interest which included the spatial extent of the saccade target (convergent remapping) [13]. We revisited these contradictory findings from a functional perspective by systematically assessing the transient consequences of retinotopic remapping on pre-saccadic sensitivity along and around the path of a saccade. We then introduced a novel neurobiologically inspired phenomenological model in which we proposed that the underlying pre-saccadic attentional and oculomotor signals manifest as temporally overlapping forces that act on retinotopic brain areas. We show that, contrary to the dominant spatial account, predictive remapping is neither a purely convergent nor a translational phenomenon but rather one which must include a translational, convergent and a centripetal component. We also demonstrate that, contrary to the dominant temporal account of predictive remapping, centripetal shifts towards the fovea precede and overlap with convergent shifts towards the peripheral region of interest, while translational shifts parallel to the saccade trajectory occurs later in time and overlap with convergent shifts.

Our study makes four principal contributions to a deeper understanding of predictive remapping. First, recipient retinotopic brain areas receive temporally overlapping inputs that align with a canonical order of pre-saccadic events (Fig 4B, Fig 7A, left panel). Second, the neural computations that underlie predictive remapping, obeys an inverse distance rule. Third, during an active vision (when not in a state of equilibrium) pRFs very likely possess putative transient eccentricity dependent elastic fields (pE∅s). It is likely that pE∅s are self-generated by the visual system and allow these receptive fields to undergo a remarkable degree of elasticity beyond their classical spatial extent. In fact, as our simulation results show, in the absence of these pE∅s orderly perception of the visual world would be dramatically distorted, as retinotopic cells would over-commit to certain loci in visual space at the expense of others. Finally, our study strongly suggests that the immediate availability of neural resources after the execution of the saccade is uniquely mediated by centripetal signals directed towards the current center of gaze. Indeed, without centripetal signals or pE∅s, attentional resources that are needed at the approximated location of the target would be is significantly delayed and in the extreme case non-existent (Fig 7B)

In conclusion, our study provides a mechanistic account of the neural computations and architecture that mediates predictive remapping which had eluded the field for decades. It also provides critical insights that will inform future neural investigations of this phenomenon specifically those concerned with uncovering the neural basis of pE∅s.

## Author contributions

EL & ASN conceptualized the project. EL collected the data. EL & XZ pre-processed the behavioural data. EL analysed the behavioural data. EL & ASN developed the computational model. EL implemented the model and performed the model simulations. EL & ASN wrote the manuscript.

## METHODS

### Predictive remapping experiment

#### Consenting human subjects

Eight subjects, all with corrected-to-normal visual acuity, provided written and informed consent prior to the study. Each subject, with the exception of two individuals (both authors), were compensated 20 USD per hour for their participation. The protocol for this study, the collection and storage of the data was approved by the Yale Ethic Review board and was in accordance with the Declaration of Helsinki.

#### Viewing distance

Subjects sat comfortably with their chin and forehead placed against a custom-built head support apparatus consisting of an adjustable chin rest, forehead support and a bite bar holder. The center of the experimental display was placed at a distance of 57cm from the eye. In order to ensure tight control over viewing distance and eye-position, subjects placed a bite bar (personalized deep-impression dental bite-bar) inside their mouth. The bite bar was then affixed to the head support apparatus for the duration of the experimental session.

#### Stimulus and presentation

Visual stimuli were presented on a gamma corrected LCD monitor using custom-designed Window software (Picto). All stimuli presented in the control and main experimental sessions, with the exception of the achromatic Gabor probes (0.5 degrees of visual angle [dva], 0° orientation, π/2 phase), were constructed in Picto. The display was driven by a NVIDIA graphics card with a color resolution of 8 bits per channel. Its spatial resolution was 1400 × 1050 pixels with a refresh rate of 85 Hz (11.76 ms per frame), and an average mean luminance of 38 cd/m2.

#### Eye-movement Tracking

The right eye (the dominant eye for all subjects) was tracked using an infrared video-based eye-tracker sampled at 1kHz (I-Scan Inc., Woburn, MA). At the beginning of each experimental session, we performed a custom-developed 9-point eye-calibration procedure in Picto. This procedure allowed for the correction of any drift in gaze at the onset of a trial. Overall, the gaze error within subjects varied from 0.25° to 1.0° with an average gaze error of 0.45° across subjects.

#### Contrast Sensitivity Function measurement

Prior to the main task, we measured the Contrast Sensitivity Function (CSF) at an eccentricity corresponding to the mean location at which probes were presented for the main task. CSF was measured using a set of Gabor stimuli contrasted against a grey background with 21 contrast levels ranging from 1 to 55%. At the onset of each trial, subjects were instructed to acquire and maintain fixation for at least 1000ms. Trails were aborted if the eye-position deviated from fixation by more than 1 dva. A Gabor stimulus with a randomly selected contrast level was then flashed for 20ms. Subjects had to indicate with a button-press whether they were able to detect the stimulus. Each contrast level was presented 10 times, along with a non-probe condition, which controlled for any potential false alarms. After the conclusion of this session, a logistic function was used to estimate a psychometric curve fitted to the data to compute a CSF. The contrast at which the subject could detect the stimuli with 50% accuracy was chosen as the probe contrast level for the main task. The CSF measurement sessions lasted approximately 70 minutes.

#### Cued saccade task

In the main experiment, subjects performed a cued saccade task (Fig 1A). Before the commencement of an experimental session, each subject took ∼15 minutes to adapt to the scotopic condition in the experimental room. At this time, a 9-point eye-position calibration procedure was performed (see above). Subjects initiated a trial by maintain gaze on a central fixation dot (subtending 0.5 dva in diameter) for 300ms. If the eye deviated from this dot by more than 1 dva, the trial was aborted. After a variable delay period of 0-300ms a saccade target (white circle) appeared at a 10° eccentric location (Fig 4A). Subjects were required to maintain fixation for another 300-600ms, at the end of which the fixation dot was extinguished, which served as the central movement cue to make a saccade towards the saccade target (Go_onset_, Fig 1B). The saccade target remained on the display for another 500ms after the central movement cue. A 2-dva radius tolerance window around the saccade target was used to detect whether the saccade landed on the target within this 500ms window. Subjects were provided visual feedback about the outcome of the trial by changing the colour of the saccade target to green for a correct saccade landing within the tolerance window or red for saccades landing outside the tolerance window. On 25% of the trials a Gabor probe (contrast set to 50% detection rate from the CSF measurement), was flashed for 20ms at a random time during the 300-600ms window between the onset of the saccade target and the central movement cue. On 50% of the trials, the probe was presented at a random time between 0-340ms after the central movement cue. To control for any false alarms no probe was flashed in the remaining 25% of trials. These three conditions were randomly interleaved across trials. Subjects indicated whether they detected the probe by a button press. Consecutive trials were separated by a 1000ms inter-trial interval. The spatial locations of the flashed probes depended on the experimental conditions outlined below. Each experimental session lasted about 70 minutes with a 10-minute break in the middle. Subjects took about 3 weeks to complete the main experiment.

#### Foveal probes condition

3 subjects (all female, *M*_*age*_= 22, SD =1.7) were recruited for the foveal condition. With the exception of subject XZ (*s*_*XZ*_, an author), each subject was completely naïve to the aim of the experiment. Saccade targets appeared at an eccentricity of 10° and at azimuth angles of 0° (for *s*_*XZ*_), 45° (for *s*_2_) or 315° (for *s*_3_) (Fig3A, left panel). In the event a flashed probe was displayed (on 75% of trials), this occurred with equal probability at one of 4 isotropic locations which were 2.5 dva from the fixation dot. Two of these locations were along the radial axis (the axis collinear with the fovea): farther away from fixation (*rad*_*fov*−*out*_), and between the fixation dot and the saccade target (*rad*_*fov*−*in*_). The other two locations were along the orthogonal (tangential) axis: counterclockwise (*tan*_*fov*−*ccw*_) or clockwise (*tan*_*fov*−*cw*_) (Fig1A, left panel).

#### Parafoveal probes condition

The same group of subjects who participated in the foveal condition took part in the parafoveal condition. The location of the saccade targets was the same as in the foveal condition. Here, the probes were presented around the midpoint (at eccentricity of 5°) between the fixation dot and the location of the saccade target (i.e., at the parafoveal retinotopic location mid-way along the saccade trajectory). Each flashed probe appeared 2.5° dva from this parafoveal midpoint along the radial-tangential axes. Specifically, a probe could either be flashed at the most inner radial parafoveal location (*rad*_*para*−*in*_) (i.e. location closest to the fovea, which overlaps with the *rad*_*fov*−*in*_ probe location in the fovea experiment), the least inner radial parafoveal location(*rad*_*para*−*out*_) (i.e. location furthest away from the fovea) or along the tangential axis through the parafoveal midpoint either counter clockwise (*tan*_*para*−*ccw*_) or clockwise (*tan*_*para*−*cw*_) (Fig4A, middle panel).

#### Peripheral probes condition

5 subjects (2 Males, 3 Females, *M*_*age*_= 25, SD =2.7) were recruited for this condition. With the exception of subject EL(*s*_*EL*_, an author), each subject was completely naïve to the aim of the experiment. Depending on the subject, the saccade target was presented at an eccentricity of 10° with azimuth angles of 270° (for *s*_1_), 315° (for *s*_2_), 0° (for *s*_3_), 45° (for *s*_4_), 90° (for *s*_*I*EAO_). In this condition a probe was flashed 2.5° dva from the saccade target along the radial-tangential axis. Specifically a flashed probe appeared at either the inner radial location (*rad*_*peri*−*in*_, the same location the *rad*_*para*−*out*_ probe was flashed), the outer radial location (*rad*_*peri*−*out*_), or along the tangential axis either counter clockwise (*tan*_*peri*−*ccw*_) or clockwise (*tan*_*peri*−*cw*_) with respect to the saccade target. (Fig4A, right panel).

### Data Analysis

#### Valid trials

Eye-position and push-button responses obtained from each subject were recorded at 1kHz and stored for further analyses. A trial in which the subject’s eye-position landed within the 2-dva tolerance window around the saccade target and within 500ms of the central movement cue was considered a valid trial. Across all subjects we collected a total of 5826 valid trials for the foveal condition, 5579 valid trials for the parafoveal condition, and 9473 valid trials for the peripheral condition.

#### Saccade onset and offset estimation

For each valid trial, we estimated the onset and offset of the saccade using a “displacement method”. Specifically, we calculated the variance in eye-position before the central movement cue and after the eye landed on the saccade target. We then estimated the lower and upper bounds of the 95% confidence intervals which accounted for any possible spurious movement which occurred before and after the saccade. The time points at which the eye-position deviated from these bounds were used to estimate the start (S_onset_) and end (S_offset)_ of the saccade.

#### Excluding trial with corrective saccades

We eliminated trials with corrective saccades (i.e., secondary saccades which compensated for under- or overshoots in the primary saccade), since such trials could potentially be associated with non-canonical sensitivity dynamics. Analysis revealed that such trials always had saccade durations (including both the primary and secondary components) greater than 50ms. We therefore eliminated trials with total saccade duration greater than 50ms, and also visually verified that such trials contained a corrective component. In the foveal condition, of the 5826 valid trials, 17 trials were eliminated for subsequent analyses (no-probe =6; *rad*_*out*−*fov*_=3; *rad*_*fov*−*in*_=3; *tan*_*fov*−*ccw*_=2; *tan*_*fov*−*ccw*_ =3). Similarly, 6 out of 5579 valid trials were eliminated in the parafoveal condition (no-probe =1; *rad*_*para*−*out*_=0; *rad*_*para*−*in*_=3; *tan*_*para*−*ccw*_=1; *tan*_*para*−*cw*_ =1), and 12 out of 9473 valid trials were eliminated in the peripheral condition (no-probe =1; *rad*_*peri*−*out*_=3; *rad*_*peri*−*in*_=3; *tan*_*peri*−*ccw*_=1; *tan*_*peri*−*cw*_ =4).

#### False alarm rate

We calculated the false alarm as the probability of a button press in trials in which no probe was presented. Subjects performed the task with very low false alarm rates in all experimental conditions: 2%, 1% and 1% respectively in the foveal, parafoveal and peripheral conditions.

#### Raw fluctuations in sensitivity

To estimate the visual sensitivity at different retinotopic locations during a rapid eye moment, we isolated the valid trials which included a flashed probe at one of the four spatial locations in the foveal, parafoveal and peripheral conditions. We first realigned the trial data to the saccade onset time (time zero) and calculated the probe presentation time with respect to saccade onset. We then calculated sensitivity, separately for the four spatial locations, by using a 25ms sliding window that was moved in 15ms increments. For each time window we calculated the fraction of trials with button pushes over the number of trials in which a probe was presented. To obtain a robust estimate of each subject’s sensitivity, we employed a 20-fold jackknife procedure in which the sensitivity was estimated from 95% of the data. This was repeated 20 times, each time leaving out 5% of the data, giving us a mean and error estimate of the sensitivity for each spatial location in each experimental condition (*rad*_*fov*−*out*_, *rad*_*fov*−*in*_, *rad*_*fov*−*tcc*_, *rad*_*fov*−*tcw*_ probes in the foveal experiment; *rad*_*para*−*out*_, *rad*_*para*−*in*_, *rad*_*para*−*tcc*_, *rad*_*para*−*tcw*_ probes in the parafoveal experiment, and the *rad*_*peri*−*out*_, *rad*_*peri*−*in*_, *rad*_*peri*−*tcc*_, *rad*_*peri*−*tcw*_ probes in the peripheral experiment).

Keeping in mind that in the foveal and parafoveal experiment, flashed probes at the *rad*_*fov*−*in*_ and the *rad*_*para*−*in*_ locations, and in the parafoveal and peripheral experiment, flashed probes at the *rad*_*para*−*out*_ and *rad*_*peri*−*in*_ locations were subtended at the same retinotopic location, we further combined the jackknife sensitivity data at these locations. In the same vein, symmetric points along the tangential location in the foveal, parafoveal and peripheral regions were also combined. Consequently, we obtained a total of seven radial and tangential sensitivity functions: four along the radial axis (*rad*_*fov*−*out*_,*rad*_*fov*−*in*_+*rad*_*para*−*in*_, *rad*_*para*−*out*_+*rad*_*peri*−*in*_, *rad*_*peri*−*out*_), three along the tangential axes (*tan*_*fov*−*ccw*+*cw*_, *tan*_*para*−*ccw*+*cw*_, *tan*_*peri*−*ccw*+*cw*_) where the x-axis represents the onset of the flashed probes from saccade onset, while the y-axis represents the sensitivity changes at a given retinotopic location. To then avoid any erroneous sensitivity results due to low sampling, these ten radial and tangential retinotopic function were truncated from 540ms prior to saccade onset to 130ms after saccade onset, a temporal window which extends from (i.) pre-planning, (ii.) eye-movement planning, *S*_*prep*_, (computed by taking the temporal difference of *S*_*onset*_ from *Go*_*onset*_, central movement cue), and (iii.) execution (*S*_*onset*_ to *S*_*offset*_) phases of the cued detection task.

#### Normalized sensitivity functions

Despite measuring the CSF at the mean probe location for each experimental condition, it is possible that there are baseline sensitivity differences across probe locations within a condition. To therefore control for any eccentric-dependent effect on the raw sensitivity functions, we normalized these functions by the average sensitivity over the initial 50ms in the data (540ms to 490ms prior to saccade onset).

#### Periodicity of sensitivity functions

To investigate the frequency components of the sensitivity data, we first de-trended the data by subtracting the mean sensitivity (Supplementary Fig 1B) and then applied a fast Fourier transform (FFT) on the detrended data. To further investigate whether the observed periodicities included a spatial component (i.e., were spatially dependent), we randomly shuffled the probe location identities across experimental trials and calculated the sensitivity function and periodicity of the shuffled data using the same procedures as above. The shuffling procedure was repeated 1500 times for each experiment. Finally, we performed a set of two-tailed paired-sample t-tests between the FFTs calculated from the detrended data and the shuffled data to determine statistical significance.

#### Retinotopic mechanics

The purpose of our computational model was to provide a general mathematical abstraction that reveals the spatiotemporal characteristics, computations and the neural architecture that actively mediates predictive remapping. Our model is agnostic to the specific neural mechanisms that underlie these processes and consists of two basic components: the retinotopic field and a force field (Fig 4).

#### Retinotopic field

We refer to the underlying architecture of our model as the retinotopic field (ϕ_r_). ϕ_r_ consists of a two-dimensional hexagonal grid of retinotopic receptive fields - RF_i_ - that tile visual space (Fig 4A, small orange circle). Each RF_i_ possesses an elastic field – pE∅s – which defines the boundary to which it can be perturbed. Consequently, each RF_i_ is characterized by three parameters: (a) *p*_*i*_, the location of the RF centre (x_i_, y_i_) at t_0_ (when there is no resultant force acting on the ϕ_r_).*p*_*i*_ determines the eccentricity - e_i_ of RF_i_. (b) s_i_, the radius of RF_i_ is proportional to e_i_. (c) The movement extent (M_max_) of RF_i_ is its pE∅ which is also proportional to e_i_.

#### Force field

Another layer within our model is the force field (ϕ_f_). ϕ_f_ exerts its influence on RF_i_ thus causing it to change its pi at t_0+n_. Specifically, ϕ_f_ includes three external forces: the centripetal force 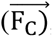, the peripheral force (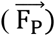), and the translational force 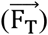and an internal force: the equilibrium force 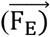. The external forces are exerted by corresponding time-varying masses: M_C_ – the mass subtended on the central region on the retina, M_P_ – the mass subtended at a peripheral site, and M_S_ – a virtual mass at infinity whose action is to produce a force in the direction of and parallel to the impending saccade. Each mass included the following distributive parts: an exponential growth, a stable plateau, and an exponential decay respectively:

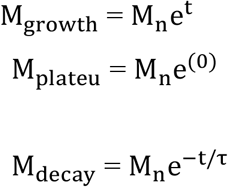

The following constitutes the possible forces experienced by RF_i_ at t_0+n_:

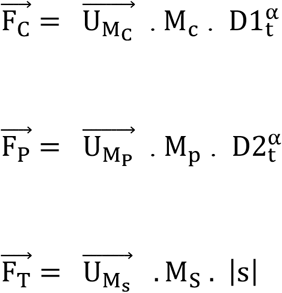

where 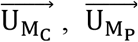, and 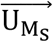 are the unit vectors in the direction of the three masses. *D*1_*t*_ and *D*2_*t*_ is the spatial difference between a RF_i_ and M_C_ or M_P_ raised to a scalar α – a distance exponent, which is the principal parameter used to control the inverse distance rule in our model. |s| is the magnitude of the impending saccadic eye movement. A resultant force 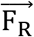, depending on the condition, could then include a single force (e.g., 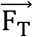), two (e.g., 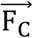 and 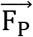), or, in the most dynamic case, all three external forces and the internal force:

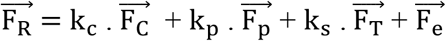

where k_*c*_ was obtained by calculating the average of 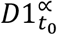 across RF_i_’s and then taking its reciprocal 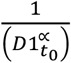. The same algorithm was used to compute k_p_, however in this case, the average of 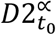 across RF_i_’s was used. It is worth noting that for a given *∝*, k_c_ and k_p_ was recalculated, but once it was obtained, it did not change at *t*_0+*n*_. Furthermore, with k_c_ and k_p_, we obtained a range of magnitude of forces for 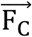 and 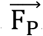. With this range, we later constructed a look-up table, where k_S_ was selected such that 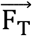 and 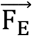 were in the same order of magnitude as 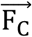 and 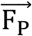. Note that as these external forces exerted their influences on RF_i_ in discrete time increments of 1ms, we calculated 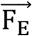 which is proportional to the displacement RF_i_ (Δd) experiences from its position at equilibrium.

#### Spatiotemporal retinotopic shifts and Density estimation

The time varying forces perturbed each RF_i_. To this end, RF_i_’s movement captured spatiotemporal retinotopic displacements. The concerted movements of the constellations of RF_i_s manifested in time-varying modulation of density at a given location of visual space. We modelled each RF_i_ as a bivariate gaussian kernel function G. To then obtain a probability density estimate, Embedded Image, which is equivalent to changes in visual sensitivity in the functional domain, we used the following equation:

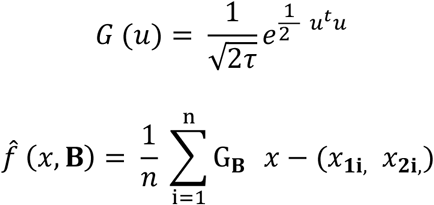

where *x*_**1i**,_ *x*_**2i**,_ denotes a sample from our bivariate distribution. B denotes the bandwidth used which was estimated using the recommendation by Bowman and Azzalini [53], with G_**B**_ as a non-negative and symmetric function (∫ *G*_*B*_(*u*)*du* = 1) defined in bivariate terms as 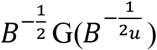. Finally, given the eccentric effects we highlighted in our behavioral data, density estimate approximate the *rad*_*fov*−*out*_, *rad*_*fov*+*para*−*in*_, *rad*_*para*+*peri*−*in*_ and *rad*_*peri*−*out*_ retinotopic locations. Further, the sensitivity we have reported were normalized by dividing density estimate by the mean of the first 110ms (i.e., the temporal window 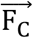 is in a steady state).

## Acknowledgments

This research was supported by NARSAD Young Investigator Grant, Ziegler Foundation Grant, Yale Orthwein Scholar Funds and Lawrence Family Young Investigator Award to ASN, and by NEI core grant for vision research P30 EY026878 to Yale University. We would like to thank Barbora Wichterlová for reading multiple edits of the manuscript.

## References

1. Gottlieb J., Oudeyer P.Y. Towards a neuroscience of active sampling and curiosity. Nat. Rev. Neurosci. 2018;19:758–770.

2. Helmholtz H.V. Dover; New York, NY: (1962). A treatise on physiological optics.

3. von Holst E, Mittelstaedt H. (1950) The reafferent principle: reciprocal effects between central nervous system and periphery. Naturwissenschaften, 37:464–476.

4. Stark L, Bridgeman B. (1983) Role of corollary discharge in space constancy. Percept. Psychophys ;34:371–380.

5. Medendorp WP. (2011). Spatial constancy mechanisms in motor control. Philos Trans R Soc Lond B Biol Sci 366: 476–491.

6. Deubel H, Bridgeman B, Schneider WX. (1998). Immediate post-saccadic information mediates space constancy. Vision Res 38: 3147–3159.

7. Duhamel, J.R., Colby, C.L., and Goldberg, M.E. (1992). The updating of the representation of visual space in parietal cortex by intended eye movements. Science, 255(5040), 90–92.

8. Walker M. F., Fitzgibbon E. J., and Goldberg M. E. (1995). Neurons in the monkey superior colliculus predict the visual result of impending saccadic eye movements. J. 746 Neurophysiol. 73, 1988–2003.

9. Sommer M. A., and Wurtz R. H. (2006). Influence of the thalamus on spatial visual processing in frontal cortex. Nature 444, 374–37710.

10. Khayat, P. S., Spekreijse, H., and Roelfsema, P. R. (2004). Correlates of transsaccadic integration in the primary visual cortex of the monkey. Proc. Natl Acad. Sci. 101, 12712–12717.

11. Nakamura, K., and Colby, C.L. (2002). Updating of the visual representation in monkey striate and extrastriate cortex during saccades. Proc Natl Acad Sci. 99(6):4026–31.

12. Tolias, A. S., Moore, T., Smirnakis, S. M., Tehovnik, E. J., Siapas, A. G., and Schiller, P. H. (2001). Eye movements modulate visual receptive fields of V4 neurons. Neuron 29(3), 757–67.

13. Zirnsak, M., Steinmetz, N. A., Noudoost, B., Xu, K. Z., and Moore, T. (2014). Visual space is compressed in prefrontal cortex before eye movements. Nature 507, 504–507.

14. Hartmann, T.S., Zirnsak, M., Marquis, M., Hamker, F.H., and Moore, T. (2017). Two types of receptive field dynamics in area V4 at the time of eye movements? Front. Syst. Neurosci. 11, 1–7.

15. Szinte, M., Jonikaitis, D., Rangelov, D., and Deubel, H. (2018). Pre-saccadic remapping relies on dynamics of spatial attention. eLife 7:e37598.

16. Zirnsak, M., Gerhards, R.G.K., Kiani, R., Lappe, M., Hamker, F.H. (2011). Anticipatory saccade target processing and the presaccadic transfer of visual features. J. Neurosci. 31(49):17887–91.

17. Rolfs, M., Jonikaitis, D., Deubel, H., and Cavanagh, P. (2011). Predictive remapping of attention across eye movements. Nat. Neurosci. 14, 252–256.

18. Jonikaitis D, Szinte M, Rolfs M, Cavanagh P. (2013). Allocation of attention across saccades. Journal of Neurophysiology. 109:1425–1434.

19. Zirnsak M, Moore T. (2014) Saccades and shifting receptive fields: anticipating consequences or selecting targets? Trends Cogn Sci.

20. Golomb, J.D. and Mazer, J.A. (2021). Visual Remapping. Annual Review of Vision Science. Vol 7.

21. Arkesteijn, K., Belopolsky, A. V., Smeets, J., Donk, M. (2019). The Limits of Predictive Remapping of Attention Across Eye Movements. Frontiers in psychology, 10, 1146.

22. Neupane, S., Guitton, D., and Pack, C. C. (2016). Two distinct types of remapping in primate cortical area V4. Nat. Commun.7:10402.

23. Neupane, S., Guitton, D., and Pack, C.C. (2016b). Dissociation of forward and convergent remapping in primate visual cortex. Curr. Biol. 26, R491–R492.

24. Nandy, A. S., & Tjan, B. S. (2012). Saccade-confounded image statistics explain visual crowding. Nature neuroscience, 15(3), 463–S2.

25. Phillips, M. H., & Edelman, J. A. (2008). The dependence of visual scanning performance on saccade, fixation, and perceptual metrics. Vision research, 48(7), 926–936.

26. Wong-Riley, M.T. (2010) Energy metabolism of the visual system. Eye Brain, 2, 99–116.

27. Sommer MA, Wurtz RH. (2006). Influence of the thalamus on spatial visual processing in frontal cortex. Nature. 444:374–7.

28. Yao, T., Ketkar, M., Treue, S., and Krishna, B. S. (2016). Visual attention is available at a task-relevant location rapidly after a saccade. eLife, 5 (e18009), 1–12.

29. Wang, X., Fung, C. A., Guan, S., Wu, S., Goldberg, M. E., and Zhang, M. (2016). Perisaccadic receptive field expansion in the lateral intraparietal area. Neuron 90, 400–409.

30. Chen CY, Hoffmann KP, Distler C, Hafed ZM. (2019) The Foveal Visual Representation of the Primate Superior Colliculus. Curr Biol. 8;29(13):2109–2119.e7.

31. Hafed, Z. M., and Clark, J. J. (2002). Microsaccades as an overt measure of covert attention shifts. Vision Res, 42(22):2533–45.

32. Hafed Z.M. (2013). Alteration of visual perception prior to microsaccades, Neuron. 77(4):775–786.

33. Sommer, M.A., Wurtz, R.H. (2002). A pathway in primate brain for internal monitoring of movements. Science 296, 1480–1482.

34. Matin, E. Saccadic suppression: a review and an analysis. (1974). Psychol Bull 81, 899–917.

35. Zuber, B. L. and Stark, L. Saccadic suppression: elevation of visual threshold associated with saccadic eye movements. (1966). Exp Neurol 16, 65–79.

36. Idrees, S., Baumann, M.P., Franke, F., Munch, T.A., and Hafed, Z.M. (2020). Perceptual saccadic suppression starts in the retina. Nat. Commun. 11, 1977.

37. Fiebelkorn, I.C., Saalmann, Y.B., and Kastner, S. (2013). Rhythmic sampling within and between objects despite sustained attention at a cued location. Current biology: CB 23, 2553–2558.

38. Benedetto, A., Morrone, M.C., (2019). Visual sensitivity and bias oscillate phase-locked to saccadic eye movements. J. Vis. 19, 15.

39. Benedetto, A., and Morrone, M. C. (2017). Saccadic suppression is embedded within extended oscillatory modulation of sensitivity. The Journal of Neuroscience, 37 (13), 3661–3670.

40. Hogendoorn, H. (2016). Voluntary saccadic eye movements ride the attentional rhythm. Journal of Cognitive Neuroscience, 28 (10), 1625–1635.

41. Bosman CA, Womelsdorf T, Desimone R, Fries P. A microsaccadic rhythm modulates gamma-band synchronization and behavior. J Neurosci., 29;29(30):9471–80. (2009).

42. Levin N, Dumoulin SO, Winawer J, Dougherty RF, Wandell BA. (2010) Cortical maps and white matter tracts following long period of visual deprivation and retinal image restoration. Neuron. 65:21–31.

43. Ross, J., Morrone, M.C., and Burr, D.C. (1997). Compression of visual space before saccades. Nature 386, 598–601.

44. Kaiser, M., and Lappe M. (2004) Perisaccadic mislocalization orthogonal to saccade direction. Neuron. 22;41(2):293–300.

45. Sperry, R.W., (1950). Neural basis of the spontaneous optokinetic response produced by visual inversion. J. Comp. Physiol. Psychol. 43, 482–489.

46. Deubel, H., Schneider, W.X. (1996) Saccade target selection and object recognition Evidence for a common attentional mechanism. Vision Res; 36:1827–1837.

47. Carrasco, M., Evert, D.L. Chang, I., Katz, S.M. (1995). The eccentricity effect: Target eccentricity affects performance on conjunction searches. Perception & Psychophysics, 57 (8) 1241–126.

48. Deubel, H. (2008) The time course of presaccadic attention shifts. Psychol Res, 72(6):630–40.

49. Melcher D, Colby CL (2008) Trans-saccadic perception. Trends Cogn Sci 12:466–473.

50. Melcher, D. (2007) Predictive remapping of visual features precedes saccadic eye movements. Nat. Neurosci. 10, 903–907.

51. Mays L.E., Sparks DL. J. (1980) Dissociation of visual and saccade-related responses in superior colliculus neurons. Neurophysiol. 43(1):207–32.

52. Goldberg ME, Bruce CJ. Primate frontal eye fields. III. (1990) Maintenance of a spatially accurate saccade signal. J Neurophysiol 64: 489–508.

53. Bowman, A. W. and Azzalini, A. (1997). Applied Smoothing Techniques for Data Analysis: the Kernel Approach with S-Plus Illustrations. Oxford University Press, Oxford.

