## Supplementary material for "Canonical retinotopic shifts under an inverse force field explain predictive remapping": Supp. Fig 1

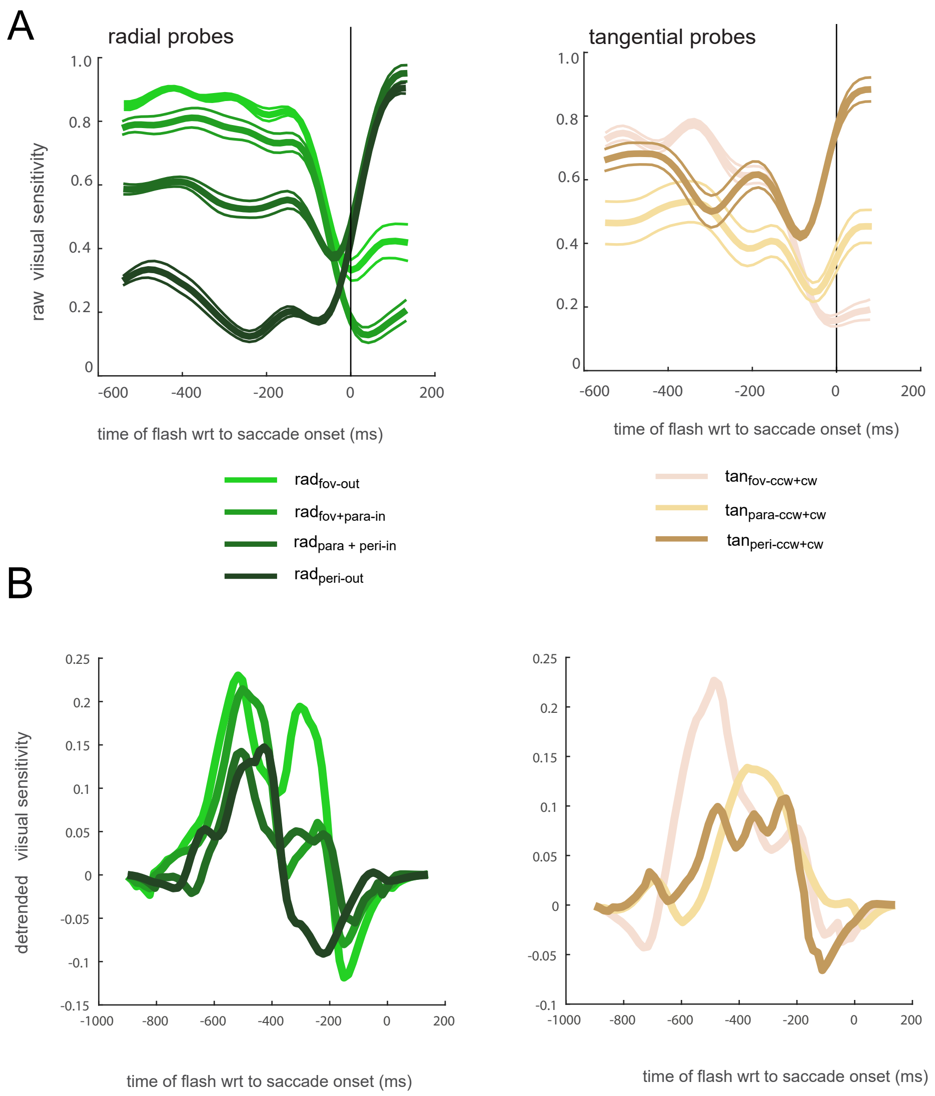


**Supplementary Fig. 1. Raw, detrended visual sensitivity**

(A) Raw sensitivity along the radial axis (left panel) and the tangential axes (right panel).

(B) Detrended sensitivity along a much wider temporal window along with the application of an Hanning window along the radial axis (left panel) and the tangential axes (right panel).
