## Supplementary material for "Canonical retinotopic shifts under an inverse force field explain predictive remapping": Supp. Fig 2

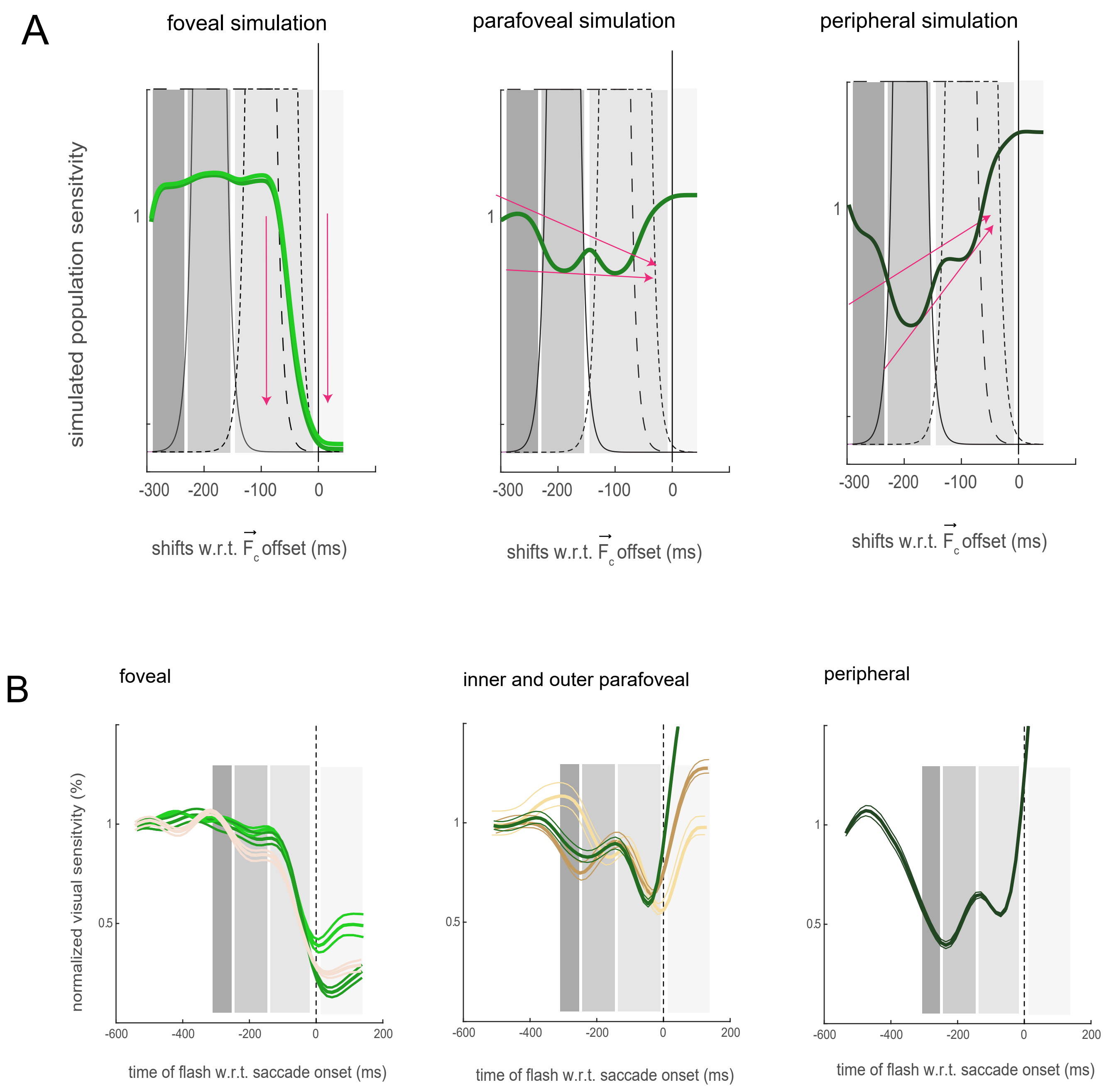


**Supplementary Fig. 2** Sensitivity functions across visual space within distinct pre and post saccadic windows.
A) Simulated population neural sensitivity. The arrows represent the general trend each sensitivity signature follows which mirrors the general signatures along radial and tangential axes observed in the behavioural experiments.see Fig.2 for comparsion.

B) Sensitivity functions across visual space from foveal, parafoveal and peripheral experiments.
